# Salidroside protects lipopolysaccharide-induced acute lung injury by regulating miR-145/ cytosolic phospholipase A_2_

**DOI:** 10.1101/2021.11.04.467383

**Authors:** Lanxin Gu, Zhaoling Shi

## Abstract

Salidroside is one of the main active components from the root of *Rhodiola rosea*. Previous reports showed that salidroside exhibits anti-inflammatory properties, but the underlying mechanisms are not fully understood. Here, we observed the effects of salidroside on lipopolysaccharide (LPS)-induced acute lung injury (ALI) both *in vivo* and *in vitro*. As revealed by survival study, salidroside reduced mortality of rats and prolonged their survival time. Meanwhile, salidroside significantly improved LPS-induced lung histopathologic changes, decreased lung wet-to-dry and lung-to-body weight ratios, inhibited lung myeloperoxidase (MPO) activity. Salidroside also suppressed the expression of cytosolic PLA_2_ (cPLA_2_), the activity of phospholipase A_2_ (PLA_2_) in LPS-treated rats and the metabolites of PLA_2_ in bronchoalveolar lavage fluid (BALF), which was confirmed by results of prostaglandin E_2_ (PGE_2_), leukotriene B_4_ (LTB_4_) and thromboxane B_2_ (TXB_2_) detection. And the expression of microRNA-145 in LPS-treated rats was up-regulated by salidroside. Besides, salidroside raised the level of miR-145and reduced PLA_2_ activity in LPS-induced A549 cells in a concentration-dependent manner, which was obviously reversed by miR-145 inhibition. In conclusion, the current study demonstrated that salidroside exhibited a protective effect on LPS-induced ALI by inhibiting of the inflammatory response, which may involve in the up-regulation of miR-145 and the suppression of cPLA_2_.

**Highlights:**

1. Salidroside reduces acute lung injury by inhibiting the increment and metabolism of phospholipase A2;
2. Salidroside inhibits LPS-induced PLA2 increase dependent on miR-145;
3. The inhibitory effect of Salidroside on Phospholipases A2 provides a link between the identification of new targets and potential new therapeutic agents for the treatment of acute lung injury.

## 1. Introduction

Acute lung injury (ALI) and acute respiratory distress syndrome (ARDS), the common and devastating clinical syndromes of acute respiratory failure in the critically ill ICU patients with high rates of morbidity and mortality[1, 2], are characterized by severe hypoxemia and uncontrolled accumulation of inflammatory cells into different compartments of the lungs, and accompanied by cytokine release and inflammatory activation of recruited or resident cells[3, 4]. Despite prosperous improvement in the treatment of ALI/ARDS, there is still a tremendous need to explore the underlying pathophysiological mechanisms of ALI/ARDS and to prevent and cure these syndromes.

Phospholipases A_2_ (PLA_2_), an ubiquitous superfamily of enzymes implicated in various inflammatory processes, catalyzes the hydrolysis of membrane phospholipids generating pro-inflammatory mediators such as thromboxanes, prostaglandins, leukotrienes and platelet-activating factor[5, 6], which are potentially involved in the development of ALI/ARDS[7, 8]. Therefore, PLA_2_ is a key enzyme for the production of these inflammatory mediators and plays an important role in ALI/ARDS. Inhibition of its activity may be a useful therapeutic strategy against inflammation.

Salidroside, an active constituent extracted from the root of *Rhodiola rosea*, exhibits many biological activities including anti-aging, anti-cancer, anti-inflammatory, anti-hypoxia and anti-oxidative properties[9–11], and has been commonly used in traditional oriental herbal medicine for diabetes, hypertension, fatigue and hypoxia[12, 13]. Previous study showed that salidroside possessed potentially beneficial anti-eicosanoid properties by suppressing the release of prostaglandin E_2_ (PGE_2_) and reducing the levels of thromboxanes B_2_ (TXB_2_) *in vitro* [13]. However, little is known about its mechanism. Additionally, the crucial effect of miRNAs on ALI/ARDS has been confirmed in recent researches[14]. It has been reported that miRNAs act as key regulators to control the process of inflammation, metabolism and repair in alveolar epithelial cells. As an important miRNA, miR-145 can be regulated by salidroside in a concentration-dependent manner in osteoarthritis injury model[15]. However, the role of salidroside and miR-145 in ALI/ARDS remains not fully investigated. In this study, therefore, we explore the effects and the underlying mechanism of salidroside on lipopolysaccharide (LPS)-induced acute lung injury both *in vivo* and *in vitro*. Results demonstrate that salidroside reduced lethality in LPS-treated rats and significantly attenuated the severity of lung injury, which was probably associated with the up-regulation of miR-145 and the inhibition of cPLA_2_.

## 2. Material and Methods

### 2.1 Animals and reagents

Adult male Sprague-Dawley rats (7–8 weeks old, and 200–250g weight) were obtained from the animal center (Fourth Military Medical University, Xi’an, and P. R. China). Rats were kept in a temperature-controlled house with 12-hour light-dark cycles. All experiments were approved by Animal Care and Use Committee at Fourth Military Medical University and were in accordance with the Declaration of the National Institutes of Health Guide for Care and Use of Laboratory Animals (Publication No.85-23, revised 1985).

Salidroside (purity is 99%, structure shown in Fig. 1A) was purchased from the National Institute for the Control of Pharmaceutical and Biological Products (NICPBP, Beijing, China). The enzyme-linked immunosorbent assay (ELISA) kits for prostaglandin E_2_ (PGE_2_) and leukotriene B_4_ (LTB_4_) were obtained from R&D Systems Inc. (Minneapolis, MN, USA). The kit for determination of myeloperoxidase (MPO) activity was obtained from Jiancheng Bioengineering Institute (Nanjing, China). Thromboxane B_2_ (TXB_2_) radioimmunoassay kit was purchased from Radioimmunoassay Institute of General Hospital of PLA (Beijing, China). Anti-cPLA_2_ and β-actin monoclonal antibodies were obtained from Santa Cruz Biotechnology Inc. (Santa Cruz, CA, USA). The EnzChek® phospholipaseA_2_ assay kit, DMEM medium and fetal bovine serums were purchased from Invitrogen Inc. (Carlsbad, CA, USA). LPS (Escherichia coli lipopolysaccharide, 055:B5), all the other reagents were obtained from Sigma Sigma-Aldrich Inc. (St. Louis, MO, USA). The purity of all chemical reagents was at least analytical grade.

**Fig.1.**
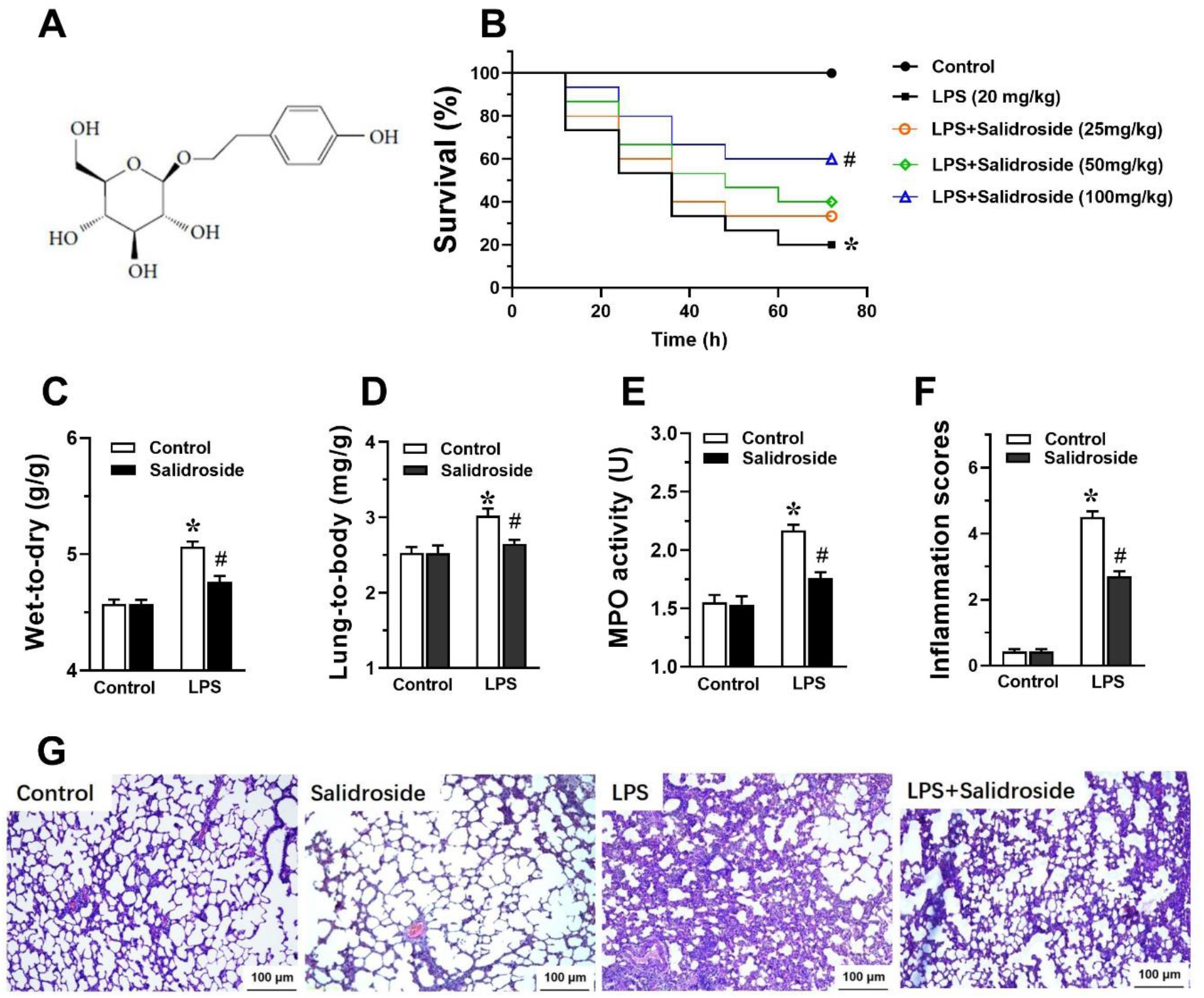
Salidroside reduced LPS-induced accumulative mortalities and improved LPS-induced lung injury in rats. (A) The chemical structure of salidroside (purity is 99%). (B) The effect of salidroside on LPS-induced accumulative mortalities. Rats were challenged by LPS (20mg/kg) with or without different doses of salidroside treatment (25, 50 or 100 mg/kg). Survival was observed for 12, 24, 36, 48, 60 and 72 h. Experiments were performed with littermate rats and each group contains 15 animals. Survival data were presented by the Kaplan Meier method and comparisons were made by the log rank test. \**P* < 0.05 vs. control group, #*P* < 0.05 vs. LPS group. Effects of salidroside on lung wet/dry weight ratios (C), lung /body weight ratios (D), myeloperoxidase (MPO) activity (E) in lung tissues, were assessed, respectively. (F-G) Histopathologic examination was performed to assess inflammation scores. The bar represents 20 μm. Data are means ±S.D., *n* = 10. \**P* < 0.05 vs. control group, #*P* < 0.05 vs. LPS group.

### 2.2 Survival studies

For the assessment of mortality rates, rats were given intraperitoneally 20 mg/kg LPS with or without different doses of salidroside (25, 50 or 100 mg/kg) treatment 0.5 h after LPS injection. The mortality of rats was recorded every 12 h for 3 days after the LPS injection. Experiments were performed with littermate rats and each group contains 15 animals.

### 2.3 Model and grouping

To further study the role of salidroside on ALI, rats were randomly divided into 4 groups, 1) Control group (*n* = 10) and 2) Salidroside group (*n* = 10): rats received saline or 100 mg/kg of salidroside intraperitoneally; 3) LPS group (*n* = 10): rats received 10 mg/kg of LPS intraperitoneally; 4) LPS+ salidroside group (*n* = 10): 100 mg/kg of salidroside was administered intraperitoneally 0.5 h after LPS administration. In all groups, measurements were made at 6 h after LPS or saline administration.

### 2.4 Histological study

At the end of experiments, the lower lobe of the right lung was fixed with 10% formalin for 24 h. After fixation, the tissues were embedded in paraffin and cut into 5 μm sections, and then stained with hematoxylin-eosin. Microscopic evaluation was performed to characterize lung injury.

### 2.5 Myeloperoxidase (MPO) activity assay and lung wet/dry ratios assessment

The upper lobe of the right lung was removed and MPO activity was measured. Briefly, the weighed lung tissue samples were frozen and homogenized in cool normal saline. The homogenates were then performed according to the manufacturer's instructions.

The remaining lung tissues were weighed immediately (wet weight), and then dried to constant weight at 50 °C for 72 h and weighed again (dry weight). The ratios of lung wet/dry weight were finally calculated to quantify the magnitude of pulmonary edema.

### 2.6 Preparation of bronchoalveolar lavage fluid (BALF) and measurements

BALF was performed (3 ml ice-cold phosphate-buffered saline three times) in all groups. In each rat, 90% (2.7ml) of the total injected volume was consistently recovered. After BALF was centrifuged at 1000g for 20 min at 4 °C, the supernatant was stored at −70 °C for subsequent measurements.

### 2.7 Enzyme-linked immunosorbent assay (ELISA) and radioimmunoassay

BALF samples were added into a 96-well plate. The concentrations of prostaglandin E_2_ (PGE_2_) and leukotriene B_4_ (LTB_4_) in BALF were determined by using commercially available ELISA kits according to the manufacturer’s instructions, respectively. These assay are based on the forward sequential competitive binding technique in which PGE_2_ or LTB_4_ presents in a sample competes with horseradish peroxidase (HRP)-labeled PGE_2_ or LTB_4_ for a limited number of binding sites on the primary polyclonal antibody. PGE_2_ or LTB_4_ in the sample is allowed to bind to the antibody in the first incubation. During the second incubation, HRP-labeled PGE_2_ or LTB_4_ binds to the remaining antibody sites. Following a wash to remove unbound materials, a substrate solution is added to the wells to determine the bound enzyme activity. The color development is stopped, and the absorbance is read at 450 nm. The intensity of the color is inversely proportional to the concentration of PGE_2_ or LTB_4_ in the sample.

The levels of thromboxane B_2_ (TXB_2_) was determined by radioimmunoassay kit according to the manufacturer’s instructions. Briefly, the BALF samples were thawed and incubated overnight at 4°C with iodine 125 labeled TXB_2_ and anti-TXB_2_ serum in a gamma globulin buffer. The next day, bound and free fractions were separated by polyethyleneglycol 6000 precipitation followed by centrifugation at 2000g at 40C for 10 min. The radioactivity of the pellet corresponding to the bound fraction was counted for 1 min in a gamma counter.

### 2.8 Western blot analysis for cPLA_2_

Total proteins in lung tissues were extracted. Protein concentrations were assayed using a coomassie brilliant blue assay. Samples were separated on a denaturing 12% SDS-polyacrylamide gel and transferred to a nitrocellulose membrane followed by incubation with primary antibodies for cPLA_2_ (1:500). Anti-β-actin antibody was used at a dilution of 1:10,000. Immunoreactivity was visualized with corresponding peroxidase-conjugated secondary antibodies and the relative content of target proteins was detected by chemiluminescence.

### 2.9 Cell culture

The human lung adenocarcinoma epithelial cell line, A549 cells obtained from American type culture collection (ATCC, Rockville, MD, USA), were maintained in DMEM medium supplemented with 10% fetal calf serum, 100 U/ml of penicillin and 100 μg/ml of streptomycin at 37 °C in a humidified atmosphere containing 5% CO_2_ and 95% air. The medium was changed every 3-4 d. The stock solutions of all the drugs were prepared in DMEM medium.

### 2.10 Methyl thiazolyl tetrazolium (MTT) assay

Cell viability was measured using the MTT assay. Briefly, A549 cells were respectively seeded into 96-well plates at 1×10^5^ cells/ml, and incubated in DMEM medium supplemented with 5% fetal calf serum for 24 h. Next, the cells were activated with 1μg/ml of LPS for 24 h in the presence or absence of salidroside (0, 0.5, 1, 2μM) for another 2 h. Then 5μl of MTT solution in PBS (5 mg/ml) was added to each well. After 4 h of incubation, MTT solution was discarded carefully, and 100μl of pure dimethyl sulfoxide was added to each well to dissolve the formazan crystals. The amount of MTT formazan was quantified spectrophotometrically by measuring the absorbance at 550 nm. Each concentration was tested in triplicate.

### 2.11 Quantitative real-time polymerase chain reaction (qRT-PCR)

qRT-PCR was used to detect the expression of miR-145 in lung tissues and A549 cells. Ttotal RNA was isolated from tissues or cells by using TRIzol reagent(Invitrogen) according to the manufacturer's protocol. Reverse transcription was performed by using cDNA Reverse Transcriptor Kit (Applied Biosystems, California, Carlsbad, USA). qRT-PCR was performed by using One Step SYBR® PrimeScript® PLUS RT-RNA PCR Kit (TaKaRa Biotechnology, Dalian, China). Relative expression levels were analyzed by using the 2^−ΔΔCT^ method.

### 2.12 miRNAs transfection

The expression plasmids of miR-145 inhibitor and its control were synthesized by Sheng Gong Co. (Shanghai, China). In brief, A549 cells were grown in 96-well plates at 1×10^5^ cells/ml, and incubated for 24h. Subsequently, miR-145 inhibitor and its control were transfected into A549 cells by using Lipofectamine 3000 reagent (Invitrogen, USA) following the manufacturer's instruction.

### 2.13 PLA_2_ activity assay in BALF and cell supernatant

Measurements of PLA_2_ activity in BALF and in A549 cell supernatants were made following the manufacturer’s protocol to investigate the underlying mechanism that salidroside generated protective effects against LPS. PLA_2_ from honey bee venom supplied with the kit was the positive control. The assay was conducted at 26°C, and fluorescence emission at 515 nm was detected after an incubation step of 10 min by a SpectraMax® M5/M5e Microplate Reader (Molecular Devices, CA, USA). The PLA_2_ activity in control (medium alone) cells was taken as 100%.

### 2.14 Statistical analysis

Data are expressed as means ± S.D., and statistical analysis was performed with analysis of variance (one-way ANOVA or two-way ANOVA), followed by a Tukey test for multiple comparisons. Survival data was presented by the Kaplan Meier method and comparisons were made by the log rank test. A statistical difference was accepted as significant if *P* < 0.05.

## 3 Results

### 3.1 Salidroside reduced LPS-induced accumulative mortalities

As shown in Fig. 1B, salidroside significantly reduced LPS-induced death. The accumulative mortalities during 3 days in 100 mg/kg of salidroside were about 60% which were observably lower than that in LPS group (about 80%, *P*<0.01). But 25mg/kg and 50mg/kg of salidroside failed to protect against death (*P*>0.05). Therefore 100mg/kg of salidroside was used to make further study.

### 3.2 Salidroside improved LPS-induced lung injury

Firstly, the lung wet/dry (W/D) weight and the lung/body weight ratios as the indexes of lung edema were detected (Fig. 1C-D). No significant difference was found between control group and salidroside group. In LPS group, the W/D weight ratios and the lung/body weight ratios were markedly increased compared with that of control group (*P*<0.01), but administration with salidroside markedly reduced the lung edema (*P*<0.05).

Secondly, we measured MPO activity to assess the neutrophil accumulation in the lung tissues. As Fig. 1E shown, LPS caused a marked increase in MPO activity compared with that of control group (*P*<0.01). Salidroside treatment obviously suppressed MPO activity induced by LPS (*P*<0.05). Similarly, salidroside also did not affect MPO activity in lung tissues of control group.

Next, we observed the pulmonary histological changes 6 h after LPS insult by microscope. Pulmonary tissue structure and alveoli in control group and salidroside group were normal. LPS instillation increased the inflammation score, caused pulmonary edema, infiltration of inflammatory cells, and alveolar damage. However, after salidroside treatment, these changes were less pronounced compared with those in LPS group (Fig. 1F-G).

### 3.3 Salidroside inhibited LPS-induced increment of PLA_2_ and its metabolites, and upregulate the level of miR-145 in rats

We further examined the repressions of miR-145 and cPLA_2_ in lung tissues, the activity of PLA_2_ and the metabolites of PLA_2_ in BALF to investigate the underlying mechanism that salidroside generated protective effects against LPS. Results showed that after LPS administration, the expression of miR-145 was significant declined, and the cPLA_2_ was markedly increased (Fig. 2 A-B, *P*<0.01), which was efficiently reversed by salidroside treatment (*P*<0.05). And PLA_2_ activity obtained from BALF in LPS-treated rats was markedly increased compared with that in control group (*P*<0.01). But, the increment of PLA_2_ activity induced by LPS was strongly inhibited by salidroside treatment ((Fig. 2 C, *P*<0.05). Similar to PLA_2_ activity, salidroside alone did not alter basal levels of PGE_2_ (Fig.2 D), LTB_4_ (Fig.4 E) and TXB_2_ (Fig.4 F) in BALF. LPS administration markedly increased the levels of these eicosanoids (*P*<0.01, respectively), whereas salidroside evidently reduced the levels of these eicosanoids induced by LPS (*P*<0.05, respectively).

**Fig.2.**
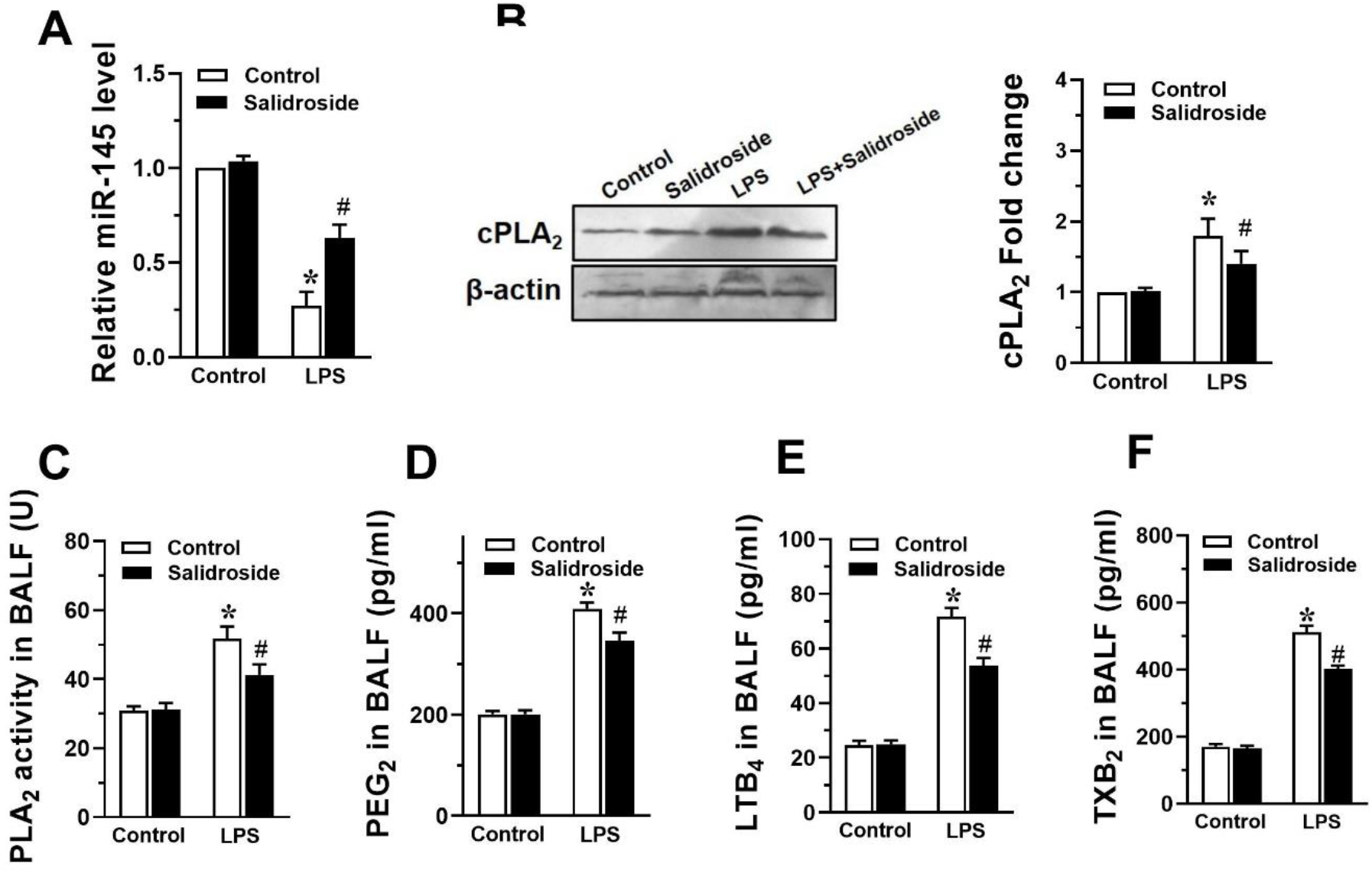
Salidroside raised the level of miR-145 and inhibited LPS-induced increment of PLA_2_ and its metabolites in BALF. The levels of miR-145 (A), the expression of cPLA_2_ (B), and the levels of PLA_2_ activity (C), prostaglandin E_2_ (PGE_2_, D), leukotriene B_4_ (LTB_4_, E) and thromboxane B_2_ (TXB_2_, F) in bronchoalveolar lavage fluid (BALF) were assessed. Data are means ±S. D., *n* = 10. \**P* < 0.05 vs. control group, ^#^*P* < 0.05 vs. LPS group.

### 3.4 Salidroside reversed LPS-induced reduction of A549 cell viability, increased the level of miR-145 and reduced LPS-induced PLA_2_ increment

MTT assays were conducted to verify the protective effects of salidroside on A549 cells. As shown in Fig. 3A, the concentration (0, 0.5, 1, 2μM) of salidroside had no effect on the viability of A549 cells. 1μg/ml of LPS markedly reduced the cell viability of A549 cells (*P*<0.01), but salidroside reversed LPS-induced reduction of the viability of A549 cells in a concentration-dependent manner (*P*<0.05).

**Fig.3.**
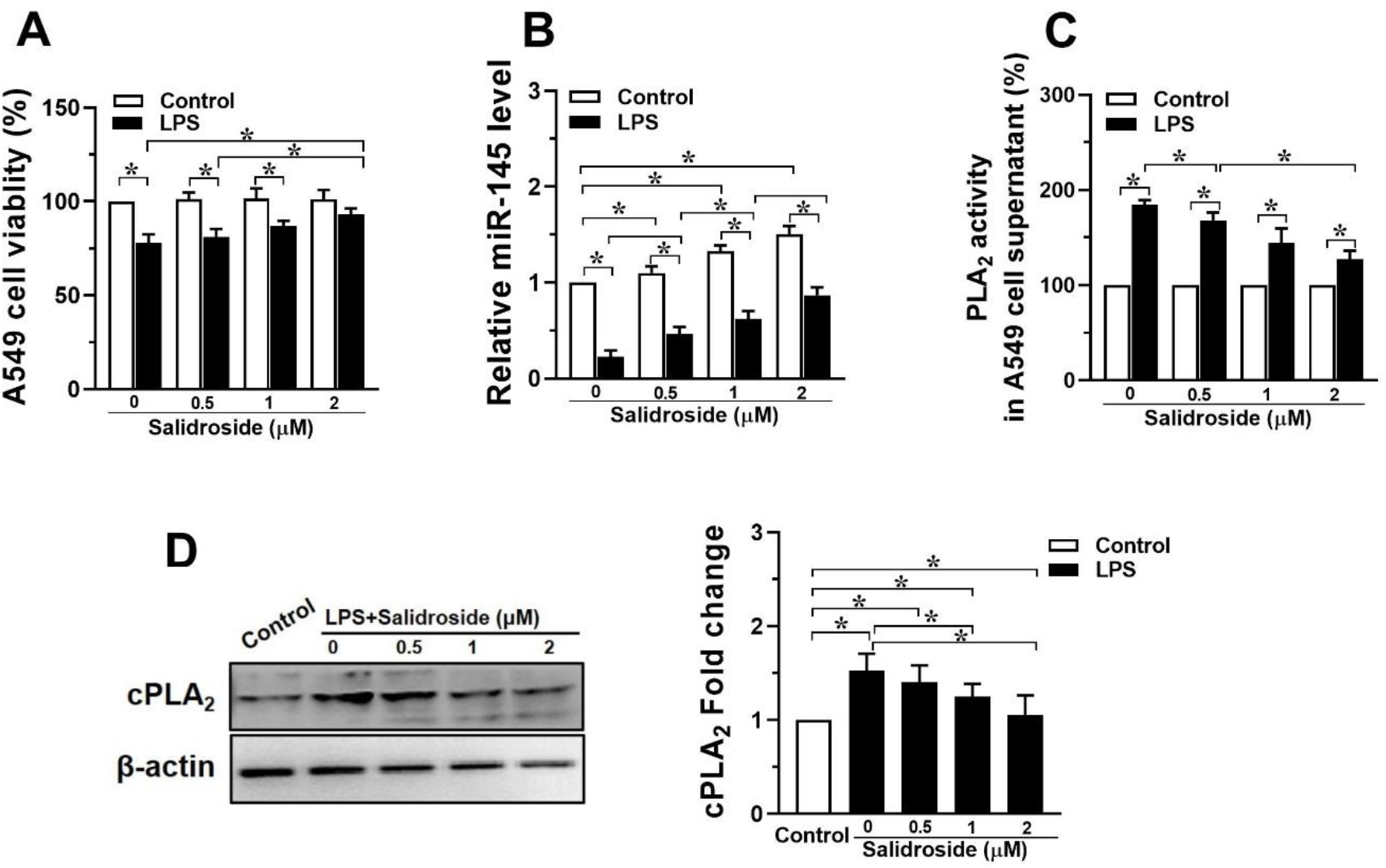
Salidroside reversed LPS-induced reduction of A549 cell viability, upregulated miR-145 level and restrained LPS-induced increment of PLA_2_. (A) A549 cells treated with different doses of salidroside (0, 0.5, 1, 2μM) after activation with LPS (1μg/ml), the viabilities were measured by MTT assay. The levels of miR-145 (B), PLA_2_ activity (C), and the expression of cPLA_2_ (D) were tested. Data are means ±S. D. from three independent experiments. \**P* < 0.05.

To clarify the relationship between LPS-induced miR-145 and salidroside, A549 cells were treated with LPS and different doses of salidroside(0, 0.5, 1, 2μM), and the expression of miR-145 was examined by qRT-PCR. As shown in Fig. 3B, salidroside promoted the expression of miR-145 in a concentration-dependent manner with or without LPS-treated (*P*<0.05). These above results suggested that miR-145 might be involved in regulation of LPS-induced inflammatory injury in A549 cells.

Additionally, the different doses of salidroside (0, 0.5, 1, 2μM) did not alter PLA_2_ activity in A549 cell supernatants, but 1μg/ml of LPS significantly increased PLA_2_ activity compared with those in control groups (Fig. 3C, *P*<0.01). Salidroside reduced LPS-induced PLA_2_ activity in a concentration-dependent manner (*P*<0.05). Similarly, the expression of cPLA_2_ induced by LPS was also suppressed (Fig. 3D, *P*<0.05)‥

### 3.5 miR-145 inhibitor blocked the effects of salidroside on LPS-induced PLA_2_ increment

Firstly, miR-145 inhibitor and its corresponding control were transfected into A549 cells to further explore the relationship of salidroside and miR-145 on LPS-induced inflammatory injury. After transfection, the level of miR-145 was significantly down-regulated, and salidroside did not raise the level of miR-145 (Fig. 4B, *P* < 0.05). Then, the effect of miR-145 inhibitor on A549 cell viability, the activity of PLA_2_ and the expression of cPLA_2_ were assessed. In Fig. 4A, the results showed that miR-145 inhibitor significantly reversed the promoting effects of salidroside on A549 cell viability (*P* < 0.05). Meanwhile, the inhibitory effects of salidroside on PLA_2_ activity and cPLA_2_ expression were also reversed by miR-145 inhibitor (Fig. 4C and D, *P* < 0.05, respectively). In a word, these results fully indicated that salidroside affected LPS induced injury by upregulation of miR-145 in A549 cells.

**Fig.4.**
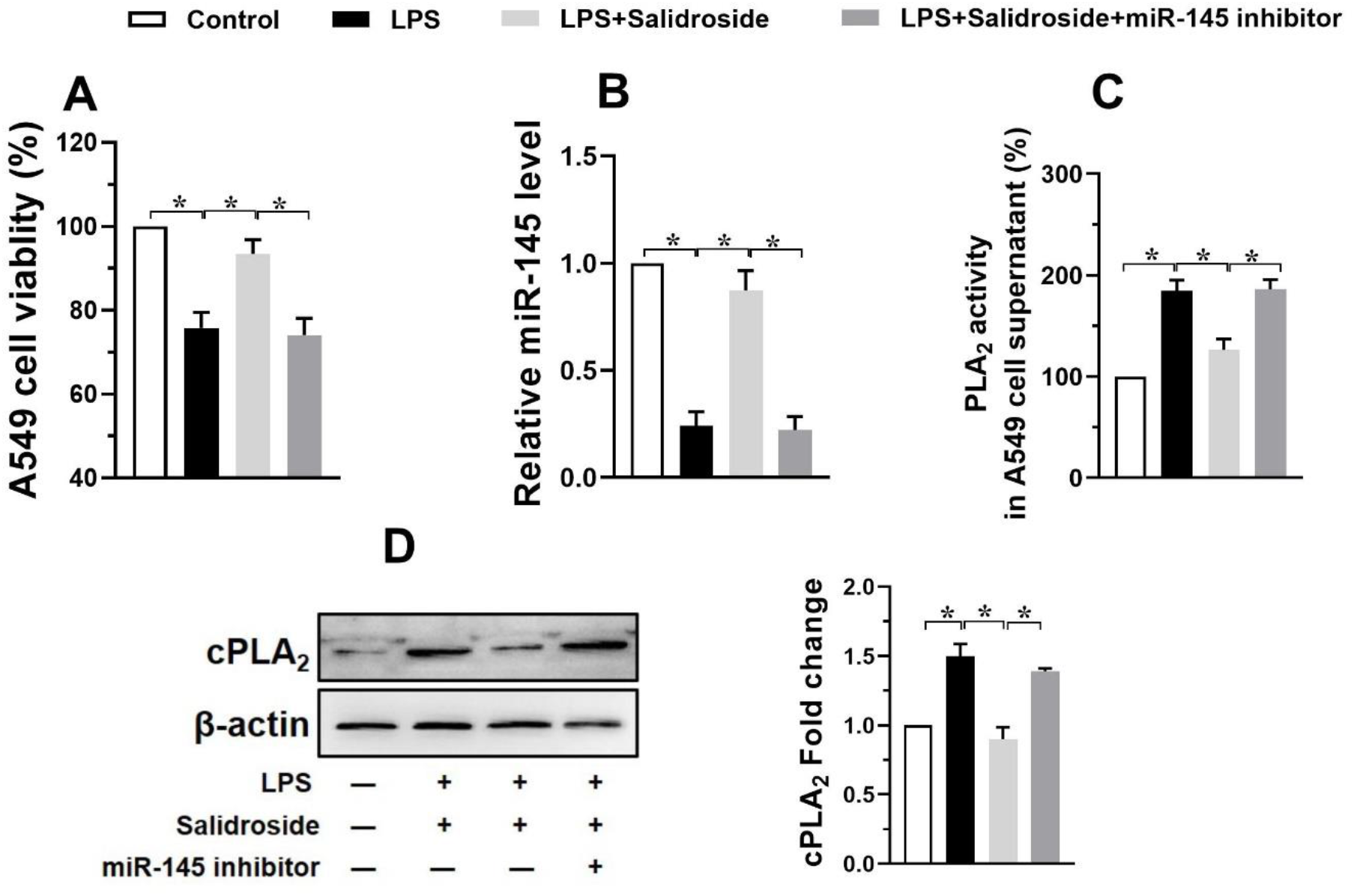
miR-145 inhibitor blocked the effects of salidroside on LPS-induced PLA2 increment. A549 cells were transfected with miR-145 inhibitor and its corresponding control to further explore the relationship of salidroside and miR-145. (A) A549 viabilities were measured by MTT assay. The levels of miR-145 (B), PLA_2_ activity (C), and the expression of cPLA_2_ (D) were also evaluated. Data are means ±S. D. from three independent experiments. \**P* < 0.05.

## 4. Discussion

In the present study, we demonstrated that salidroside played an important role in the protection of LPS-induced ALI. Results showed that salidroside could reduce mortality of rats and prolong their survival time, attenuate lung injury by inhibiting PLA_2_ activity, suppressing the expression of cPLA_2_, and cutting down the generation of pro-inflammatory eicosanoids, which was probably associated with the up-regulation of miR-145 and the inhibition of cPLA_2_, and indicated therapeutic potential for salidroside in the treatment of ALI.

*Rhodiola rosea (R. rosea)* is known as a golden or arctic root and belongs to the plant family of Crassulaceae, subfamily of Sedoideae, and genus Rhodiola.[13] It is clinically used in China or many Asian countries either alone or in combination with other herbal ingredients to prevent or manage many diseases[10, 11, 13]. Salidroside, an active constituent extracted from *Rhodiola rosea*, have the most critical therapeutic activities. It can provide a protective effect on epirubicin-induced early left ventricular regional systolic dysfunction for breast cancer patients by its antioxidant properties[10]. Salidroside also possesses antiviral activities against coxsackievirus B3 (CVB3) by adjusting antioxidant defense and regulating cytokine expression[9]. Salidroside is viewed as a promising neuroprotective drug for Alzheimer’s disease (AD) because it modulated oxidative stress and inflammatory mediators[9]. Moreover, salidroside can inhibit clinorotation-induced apoptosis in pulmonary microvascular endothelial cells through PI3K/AKT pathway [16]. In particular, the anti-inflammatory effects of salidroside were investigated on LPS-induced ALI in mice and LPS-stimulated RAW 264.7 macrophages[17]. Results shown that salidroside increased mouse survival, alleviated the production of inflammatory cytokines including tumor necrosis factor-α (TNF-α), interleukin-6 (IL-6) and interleukin-1β (IL-1β), and blocked the activation of NF-қB and ERK/MAPKs signalling pathways[18]. These results may provide a potential mechanism that explains the anti-inflammatory and antioxidant activities of salidroside and suggest a possible use of salidroside in the treatment of inflammatory diseases.

In the present study, we found that salidroside significantly reduced mortality, prolonged survival time of rats, attenuated the severity of ALI and relieved pulmonary edema and vascular permeability. The histopathologic results showed that there were great improvements after salidroside treatment. The increased wet-to-dry ratios, lung-to body weight ratios, MPO activity were markedly reduced by treatment of salidroside. Additionally, 2μM of salidroside protected A549 cells from LPS-induced injury as revealed by MTT assay. All above results suggested the potential role of salidroside in improvement of lung injury.

Furthermore, we investigated the possible mechanism of salidroside protective effects against ALI. It is well known that PLA_2_ one of the important mediators contributing to ALI/ARDS. PLA_2_ disturbs pulmonary function by hydrolyzing lung surfactant phospholipids and produces a large amount of pro-inflammatory arachidonic acids[6], which leads to the representative pathological changes of ALI/ARDS characterized by the increased alveolar-capillary barrier permeability and lung oedema formation[7]. PLA_2_s are now segregated into six major classes based on biochemical properties: secretory PLA_2_s (sPLA_2_s), cytoplasmic PLA_2_s (cPLA_2_s), calcium-independent PLA_2_s (iPLA_2_s), lysosomal PLA_2_s, platelet-activating factor acetylhydrolases (PAF-AHs), and PLA_2_s of bacterial origin[5]. Almost all of these PLA_2_ isoforms have been reported to contribute to the process of lung infection and inflammation. Administration of PLA_2_ inhibitors, such as S-5920/LY315920Na and arachidonyl trifluoromethyl ketone could attenuate LPS-induced ALI[7]. Consequently, modulating the PLA_2_ activity may represent a therapeutic approach to ALI/ARDS. In the current study, we found that LPS aggrandized PLA_2_ activity both in BALF and in the supernatants of A549 cell, increased cPLA_2_ expression in the lung tissues, whereas salidroside could down-regulate the increased PLA_2_ activity and cPLA_2_ expression induced by LPS. Moreover, salidroside reduced the release of PGE_2_, LTB_4_ and TXB_2_, which further confirms that salidroside inhibited LPS-induced PLA_2_ activity.

MicroRNAs are small non-coding RNA that plays a crucial role in many disease processes, including malignancy, inflammation, metabolism and repair processes. Abnormal expression miRNAs, such as miR-27, miR-377, miR-30b and miR-218 have been found in ALI[14, 19]. MiR-145 is an important molecular marker, which has been proven to mediate lung ischemia/reperfusion injury[20]. And Xu Qi et al. displayed that cPLA_2_ contributed to cerebral infarction is a target of miR-145[21]. These studies demonstrated that miR-145 may be implicated in cPLA_2_ regulation in ALI. However, whether salidroside attenuated LPS-induced ALI both *in vivo* and *in vitro* was involved in miR-145 and cPLA_2_ remain unknown. In our study, we found that the expression of miR-145 was up-regulated by salidroside, and miR-145 inhibitor significantly alleviated the protective effect of salidroside on LPS injured A549 cells.

There were still limitations of this study. First, determining the particular role of PLA_2_ on ALI/ARDS has proven quite challenging, because this enzyme represents a family of over 20 distinct proteins with various structural and biochemical characteristics. So what kind of PLA_2_ isoforms involved in the protective effect of salidroside on LPS-induced ALI remains to be further elucidated. We will further investigate the core mechanism responsible for the anti-inflammatory effects of salidroside by using specific PLA_2_ inhibitors or other methods. Second, it is also important to reiterate the relationship of salidroside, miR-145 and the individual PLA_2_ enzymes, which also needs to be tapped in further studies.

## 5 Conclusion

Although it requires further investigation, our results suggested that treatment with salidroside could significantly reduce lethality in LPS-treated rats, attenuated the severity of LPS-induced lung injury both *in vivo* and *in vitro*. The protection of salidroside was probably associated with the up-regulation of miR-145 and the inhibition of cPLA_2_. Conclusively, the present study partially explained the anti-inflammation capacity of salidroside. It may be considered as a potential agent in treatment for ALI/ARDS.

## Acknowledgements

This work was supported by Natural Science Foundation of Shaanxi Province of China (2019JQ-544).

## References

[1] Y. Butt, A. Kurdowska, T.C. Allen, Acute Lung Injury: A Clinical and Molecular Review, Arch Pathol Lab Med, 140 (2016) 345–350.

[2] S. Spadaro, M. Park, C. Turrini, T. Tunstall, R. Thwaites, T. Mauri, R. Ragazzi, P. Ruggeri, T.T. Hansel, G. Caramori, C.A. Volta, Biomarkers for Acute Respiratory Distress syndrome and prospects for personalised medicine, J Inflamm (Lond), 16 (2019) 1.

[3] S.A. Kuldanek, M. Kelher, C.C. Silliman, Risk factors, management and prevention of transfusion-related acute lung injury: a comprehensive update, Expert Rev Hematol, 12 (2019) 773–785.

[4] M. Camprubi-Rimblas, N. Tantinya, J. Bringue, R. Guillamat-Prats, A. Artigas, Anticoagulant therapy in acute respiratory distress syndrome, Ann Transl Med, 6 (2018) 36.

[5] A. Aloulou, R. Rahier, Y. Arhab, A. Noiriel, A. Abousalham, Phospholipases: An Overview, Methods Mol Biol, 1835 (2018) 69–105.

[6] M. Murakami, H. Sato, Y. Taketomi, Updating Phospholipase A2 Biology, Biomolecules, 10 (2020).

[7] E. Kitsiouli, M. Tenopoulou, S. Papadopoulos, M.E. Lekka, Phospholipases A2 as biomarkers in ARDS, Biomed J, (2021).

[8] S.Y. Filkin, A.V. Lipkin, A.N. Fedorov, Phospholipase Superfamily: Structure, Functions, and Biotechnological Applications, Biochemistry (Mosc), 85 (2020) S177–S195.

[9] X.L. Bai, X.L. Deng, G.J. Wu, W.J. Li, S. Jin, Rhodiola and salidroside in the treatment of metabolic disorders, Mini Rev Med Chem, 19 (2019) 1611–1626.

[10] S. Sun, Q. Tuo, D. Li, X. Wang, X. Li, Y. Zhang, G. Zhao, F. Lin, Antioxidant Effects of Salidroside in the Cardiovascular System, Evid Based Complement Alternat Med, 2020 (2020) 9568647.

[11] F. Fan, L. Yang, R. Li, X. Zou, N. Li, X. Meng, Y. Zhang, X. Wang, Salidroside as a potential neuroprotective agent for ischemic stroke: a review of sources, pharmacokinetics, mechanism and safety, Biomed Pharmacother, 129 (2020) 110458.

[12] Z. Zhong, J. Han, J. Zhang, Q. Xiao, J. Hu, L. Chen, Pharmacological activities, mechanisms of action, and safety of salidroside in the central nervous system, Drug Des Devel Ther, 12 (2018) 1479–1489.

[13] W.L. Pu, M.Y. Zhang, R.Y. Bai, L.K. Sun, W.H. Li, Y.L. Yu, Y. Zhang, L. Song, Z.X. Wang, Y.F. Peng, H. Shi, K. Zhou, T.X. Li, Anti-inflammatory effects of Rhodiola rosea L.: A review, Biomed Pharmacother, 121 (2020) 109552.

[14] S. Rajasekaran, D. Pattarayan, P. Rajaguru, P.S. Sudhakar Gandhi, R.K. Thimmulappa, MicroRNA Regulation of Acute Lung Injury and Acute Respiratory Distress Syndrome, J Cell Physiol, 231 (2016) 2097–2106.

[15] Y. Han, S.C. Mu, J.L. Wang, W. Wei, M. Zhu, S.L. Du, M. Min, Y.J. Xu, Z.J. Song, C.Y. Tong, MicroRNA-145 plays a role in mitochondrial dysfunction in alveolar epithelial cells in lipopolysaccharide-induced acute respiratory distress syndrome, World J Emerg Med, 12 (2021) 54–60.

[16] Y. Wang, C.F. Xu, Y.J. Liu, Y.F. Mao, Z. Lv, S.Y. Li, X.Y. Zhu, L. Jiang, Salidroside Attenuates Ventilation Induced Lung Injury via SIRT1-Dependent Inhibition of NLRP3 Inflammasome, Cell Physiol Biochem, 42 (2017) 34–43.

[17] L. Jingyan, G. Yujuan, Y. Yiming, Z. Lingpeng, Y. Tianhua, M. Mingxing, Salidroside Attenuates LPS-Induced Acute Lung Injury in Rats, Inflammation, 40 (2017) 1520–1531.

[18] K.C. Lan, S.C. Chao, H.Y. Wu, C.L. Chiang, C.C. Wang, S.H. Liu, T.I. Weng, Salidroside ameliorates sepsis-induced acute lung injury and mortality via downregulating NF-kappaB and HMGB1 pathways through the upregulation of SIRT1, Sci Rep, 7 (2017) 12026.

[19] L.K. Lee, L. Medzikovic, M. Eghbali, H.K. Eltzschig, X. Yuan, The Role of MicroRNAs in Acute Respiratory Distress Syndrome and Sepsis, From Targets to Therapies: A Narrative Review, Anesth Analg, 131 (2020) 1471–1484.

[20] S.H. Dai, L.J. Chen, W.H. Qi, C.L. Ye, G.W. Zou, W.C. Liu, B.T. Yu, J. Tang, microRNA-145 Inhibition Upregulates SIRT1 and Attenuates Autophagy in a Mouse Model of Lung Ischemia/Reperfusion Injury via NF-kappaB-dependent Beclin 1, Transplantation, 105 (2021) 529–539.

[21] X. Qi, M. Shao, H. Sun, Y. Shen, D. Meng, W. Huo, Long non-coding RNA SNHG14 promotes microglia activation by regulating miR-145-5p/PLA2G4A in cerebral infarction, Neuroscience, 348 (2017) 98–106.

